# LCN2 promotes focal adhesion formation and invasion by stimulating Src activation

**DOI:** 10.1101/2024.06.04.597324

**Authors:** Bhagya Shree Choudhary, Nazia Chaudhary, Aditi Vijan, Dibita Mandal, Leena Pilankar, Shubham Gawand, Bushra K. Khan, Anusha Shivashankar, Rinki Doloi, Neha Joshi, Manjula Das, Sorab N. Dalal

**Author notes:** Thoracic Head and Neck Molecular Oncology, M.D. Anderson Cancer Centre, University of Texas, Houston, USA. Faculty of Life Sciences, University of Vienna, 1030 Vienna, Austria (Rinki Doloi). School of Pharmacy and Biomolecular Sciences (PBS), Royal College of Surgeons in Ireland, Dublin 2, Ireland.

## Abstract

Previous work has demonstrated that Lipocalin2 (LCN2) expression promotes invasion and migration in multiple tumor types. The mechanisms by which LCN2 promotes invasion and migration remain unclear. Previous work from our laboratory demonstrated that LCN2 promotes actin filament formation by inhibiting actin glutathionylation. In this report, we demonstrate that in addition to inhibiting actin glutathionylation, LCN2 stimulates invasion by promoting the formation of focal adhesions, which is independent of the ability of LCN2 to bind iron. LCN2 promotes focal adhesion formation by promoting the activation of c-Src by stimulating the expression of the transcription factor ETS1. ETS1 activates the expression of the protein phosphatase, PTP1B, resulting in the auto-activation of c-Src and increased paxillin phosphorylation leading to focal adhesion formation. These results demonstrate that LCN2 has iron-dependent and independent functions in promoting invasion and highlight the multiple mechanisms by which LCN2 promotes invasion and suggest that c-Src inhibitors could be used to treat invasive colorectal cancer.

**SUMMARY:** Expression of LCN2 is elevated in invasive colorectal cancer. We demonstrate that LCN2 promotes invasion by stimulating the formation of focal adhesions by promoting Src activation.

## Introduction

An increase in invasion and migration is observed in late-stage tumor cells, often leading to metastatic disease (reviewed in (Chaffer et al., 2016; Lambert et al., 2017; Valastyan and Weinberg, 2011)). The increase in invasion and migration depends on actin polymerization and the formation of focal adhesions (reviewed in (Albiges-Rizo et al., 2009; Blanchoin et al., 2014; Case and Waterman, 2015)). While various factors stimulate focal adhesion formation (Albiges-Rizo et al., 2009; Case and Waterman, 2015), many of these processes are conserved in normal cells, making them inappropriate targets for therapeutic intervention. Hence it is essential to study the processes regulating invasion and migration in tumor cells to identify therapeutic strategies for the treatment of invasive late-stage tumors.

In response to extracellular stimuli, actin filaments drive the formation of cellular protrusions called lamellipodia and filopodia (Le Clainche and Carlier, 2008). The leading edge of the cell interacts with the extracellular matrix (ECM) by forming focal adhesions that are connected to the intracellular lamellipodial actin network (Critchley, 2009). Focal adhesions assemble contractile actomyosin cables to generate traction force (Ciobanasu et al., 2012). The traction force generated by the contraction of stress fibers leads to the disassembly of FAs and retracts the trailing edge (Ridley et al., 2003). Interaction of cells with the surrounding extracellular matrix (ECM) stimulates integrin clustering which promotes autophosphorylation of focal adhesion kinase (FAK) at Y397 to relieve auto-inhibition (Schoenwaelder and Burridge, 1999). This phosphorylation creates a docking site for activated c-Src. The active FAK-Src complex phosphorylates paxillin at Y118 resulting in focal adhesion formation (Parsons et al., 2008). c-Src is inactivated by phosphorylation at Y527 leading to inhibition of kinase activity. Y527 is dephosphorylated by the protein phosphatase, PTP1B (Zhu et al., 2007), leading to autophosphorylation at Y416 and activation of c-Src (Xu et al., 1999a).

Lipocalin2 (LCN2), also known as NGAL (neutrophil gelatinase associated lipocalin), is a secreted glycoprotein (Chakraborty et al., 2012; Schmidt-Ott et al., 2006) and is required to maintain the integrity of the gastrointestinal mucosa (Playford et al., 2006). LCN2 forms a complex with bacterial and human siderophores (reviewed in (Chakraborty et al., 2012)) thereby inhibiting bacterial growth and regulating iron homeostasis in mammalian cells. LCN2 bound to iron is imported into cells by complex formation with the LCN2 receptors, such as SLC22A17, MCR4, LRP2 or Megalin, and others (Chakraborty et al., 2012; Devireddy et al., 2005; Schroder et al., 2023). The import of iron into cells by LCN2 is required for LCN2 functions in maintaining kidney homeostasis (Mori et al., 2005; Yang et al., 2002) and in regulating apoptotic cell death (Devireddy et al., 2005; Devireddy et al., 2001). LCN2 binds to and protects the matrix-metalloprotease MMP9 from auto-degradation with a concomitant increase in MMP9 activity (Yan et al., 2001). This is consistent with the observation that LCN2 can promote invasion and angiogenesis and is associated with metastasis in multiple tumor types (Chung et al., 2016; Guo et al., 2016; Lee et al., 2006; Oren et al., 2016; Tong et al., 2008).

Previous work from the laboratory demonstrated that loss of the desmosomal plaque protein Plakophilin3 (PKP3), led to an increase in migration, invasion, and tumor formation (Basu et al., 2018; Kundu et al., 2008). These phenotypes were dependent on the increased expression of LCN2 in the PKP3 knockdown clones (Basu et al., 2018). Further, additional data from the laboratory demonstrated that LCN2 could promote tumor progression and therapy resistance in colorectal cancer and could promote invasion in colorectal cancer cell lines (Chaudhary et al., 2021; Choudhary et al., 2023). The increase in invasion was due to the ability of LCN2 to inhibit the glutathionylation of actin by regulating the levels of intracellular iron and ROS. However, neutralizing ROS and chelating iron did not completely restore invasion in LCN2 low cells suggesting that LCN2 could stimulate invasion via other mechanisms.

The results in this report suggest that in addition to inhibiting actin glutathionylation, LCN2 expression leads to increased focal adhesion formation. The increase in focal adhesion formation is not dependent on the ability of LCN2 to bind iron but is dependent on the LCN2-mediated activation of the tyrosine kinase c-Src, suggesting that inhibiting Src function in invasive colorectal cancer could be a potential therapeutic mechanism in tumors with high levels of LCN2.

## Results

Previous results from the lab have demonstrated that LCN2 expression can promote invasion by inhibiting the glutathionylation of actin thus promoting actin polymerization (Choudhary et al., 2023). Since the process of invasion requires the formation of focal adhesions, we wished to determine whether LCN2 promoted focal adhesion formation. We tested this in cells with low levels of LCN2 (HCT116) where LCN2 was over-expressed and in cells with high levels of LCN2 (DLD1) where we inhibited LCN2 expression using vector-driven RNAi (SF1A-B and (Choudhary et al., 2023)). These cells were stained with antibodies to paxillin to identify focal adhesions and counterstained with fluorescent phalloidin. As shown in Fig. 1A-B, the LCN2 over-expressing cells showed increased numbers of paxillin foci as compared to the vector control. The area and perimeter of the paxillin foci were increased in the LCN2 high cells as compared to the vector control (Fig. 1C-D). In contrast, inhibiting LCN2 expression resulted in a decrease in the number, area, and perimeter of paxillin foci (Fig. 1E-H). Similarly, inhibiting LCN2 function with the anti-LCN2 monoclonal antibody, 3D12B2 (Chaudhary et al., 2021; Choudhary et al., 2023), inhibited the formation of paxillin foci (SF1C) and resulted in a decrease in the number, area and perimeter of paxillin foci in LCN2 over-expressing cells as compared to untreated cells and cells treated with a non-specific mouse IgG (MIgG) (Fig. 1I-K). These results suggested that LCN2 could promote focal adhesion formation.

**Figure 1.**
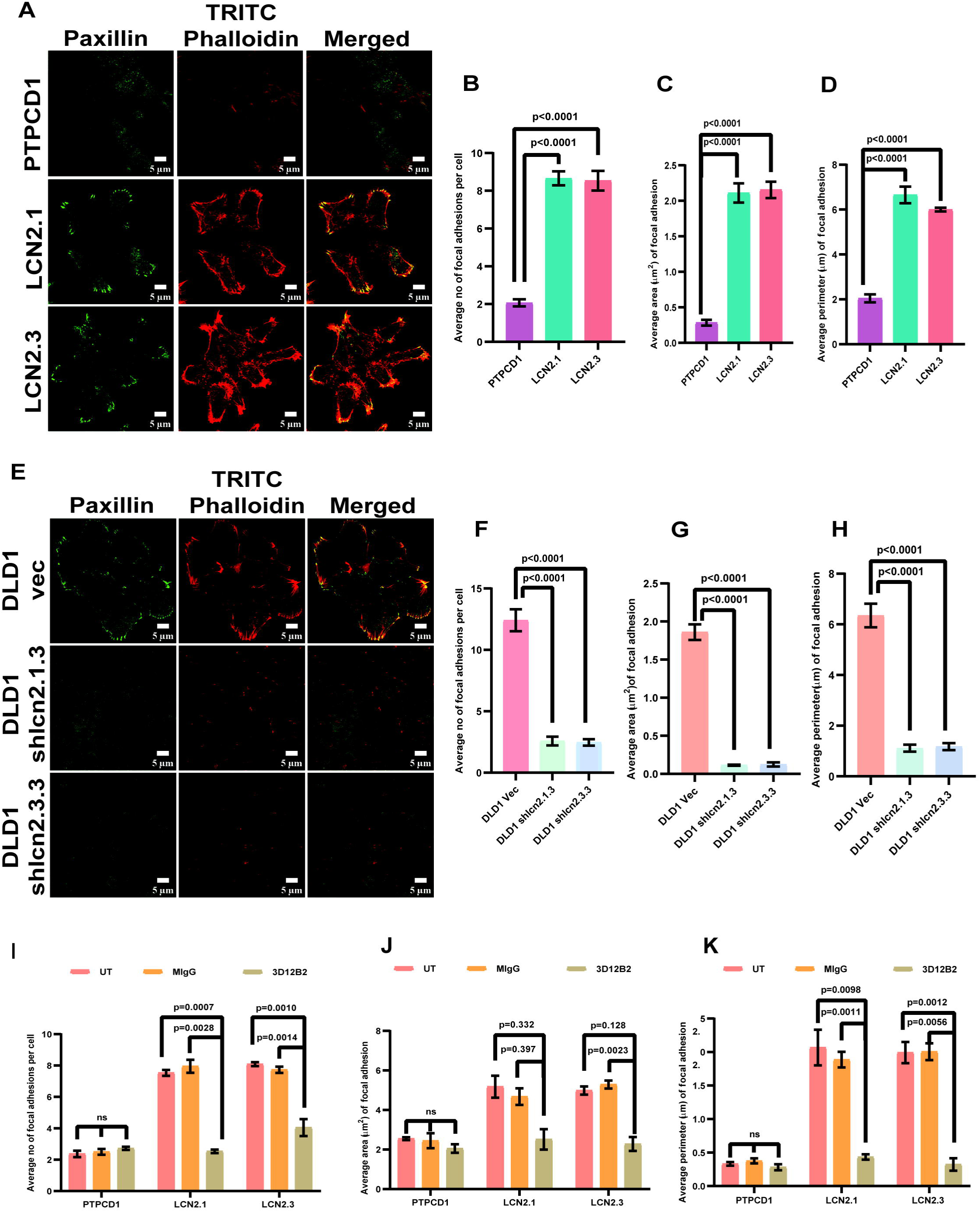
LCN2 stimulates focal adhesion formation. **A-H.** The indicated cell lines were stained with antibodies to paxillin (green) or TRITC-phalloidin (red) and imaged by confocal microscopy. Representative images are shown (A&E). The number (B&F), area (C&G), and perimeter (D&H) of focal adhesions were quantitated in three independent experiments and the mean and standard deviation were plotted. **I-K.** The indicated cell lines were untreated (UT) or treated with a non-specific mouse IgG (MIgG) or the anti-LCN2 antibody (3D12B2) and stained with antibodies to paxillin and TRITC-phalloidin. The number (I), area (J), and perimeter (K) of focal adhesions were determined in three independent experiments and the mean and standard deviation were plotted. Where indicated, p values were determined using a student’s t-test. Scale = 5µm.

The ability of LCN2 to bind to iron and regulate intracellular iron levels was required for invasion stimulated by LCN2 (Chaudhary et al., 2021; Choudhary et al., 2023). To determine if the ability of LCN2 to form a complex with iron was required for focal adhesion formation, the DLD1-derived vector control and knockdown cells were either untreated or treated with recombinant WTLCN2, Apo-LCN2 (not bound to iron) or an iron-binding defective mutant of LCN2 (K125AK134A (Bao et al., 2010)) and focal adhesion formation determined. Treatment with the WTLCN2, Apo-LCN2, and the K125AK134A mutant restored focal adhesion formation in the LCN2 knockdown clones in contrast to the untreated cells (Fig. 2A-C and SF2). As our previous data suggested that the ability of LCN2 to bind iron was required to prevent the glutathionylation of actin resulting in increased actin polymerization and invasion, we hypothesized that an actin mutant that could not be glutathionylated would promote invasion in cells treated with the iron-binding defective mutant of LCN2. Previous work has demonstrated that the major site of glutathionylation in actin is a Cysteine residue at position 374 (C374) (Wang et al., 2001). We altered this residue to Serine (C374S) and tested the ability of this mutant to restore invasion in HCT116 cells treated with WT LCN2 or the K125AK134A mutant of LCN2. As a first step, we attempted to determine the level of glutathionylation on the C374S mutant. HCT116 cells were transfected with the vector control, GFP-actinWT or GFP-actin C374S, and the extracts were incubated with the purified His-tagged proteins and the reactions purified on Nickle beads. His LCN2 served as a negative control in these assays and was present at the same levels as His GST (SF3B). HCT116 cells transfected with the vector control showed the presence of endogenous glutathionylated actin in the His GST pulldown in untreated cells and cells incubated with K125AK134A. In contrast, incubation with recombinant WT LCN2 resulted in a decrease in the levels of glutathionylated actin. The C374S mutant was expressed at levels similar to WT actin (SF3B and Fig. 2D), however, no significant decrease in glutathionylation in this mutant was observed in GST pulldown assays (Fig. 2D). This could be because actin has multiple Cysteine residues that could be modified by ROS (reviewed in (Wilson et al., 2016)). However, the actin C374S mutant could form actin filaments in cells treated with the iron-binding defective LCN2 mutant in contrast to WT actin as demonstrated by phalloidin staining (Fig. 2E and SF3C) as previously reported (Stournaras et al., 1990; Wang et al., 2001). In invasion assays, HCT116 cells transfected with WT actin showed a significant increase in invasion upon treatment with WT but not the mutant LCN2. In contrast, cells transfected with the C374S mutant showed increased invasion in cells treated with WT and mutant LCN2 (Fig. 2F and SF3D) suggesting that an actin mutant that isn’t glutathionylated at C374 can promote invasion in cells treated with the iron-binding defective mutant of LCN2.

**Figure 2.**
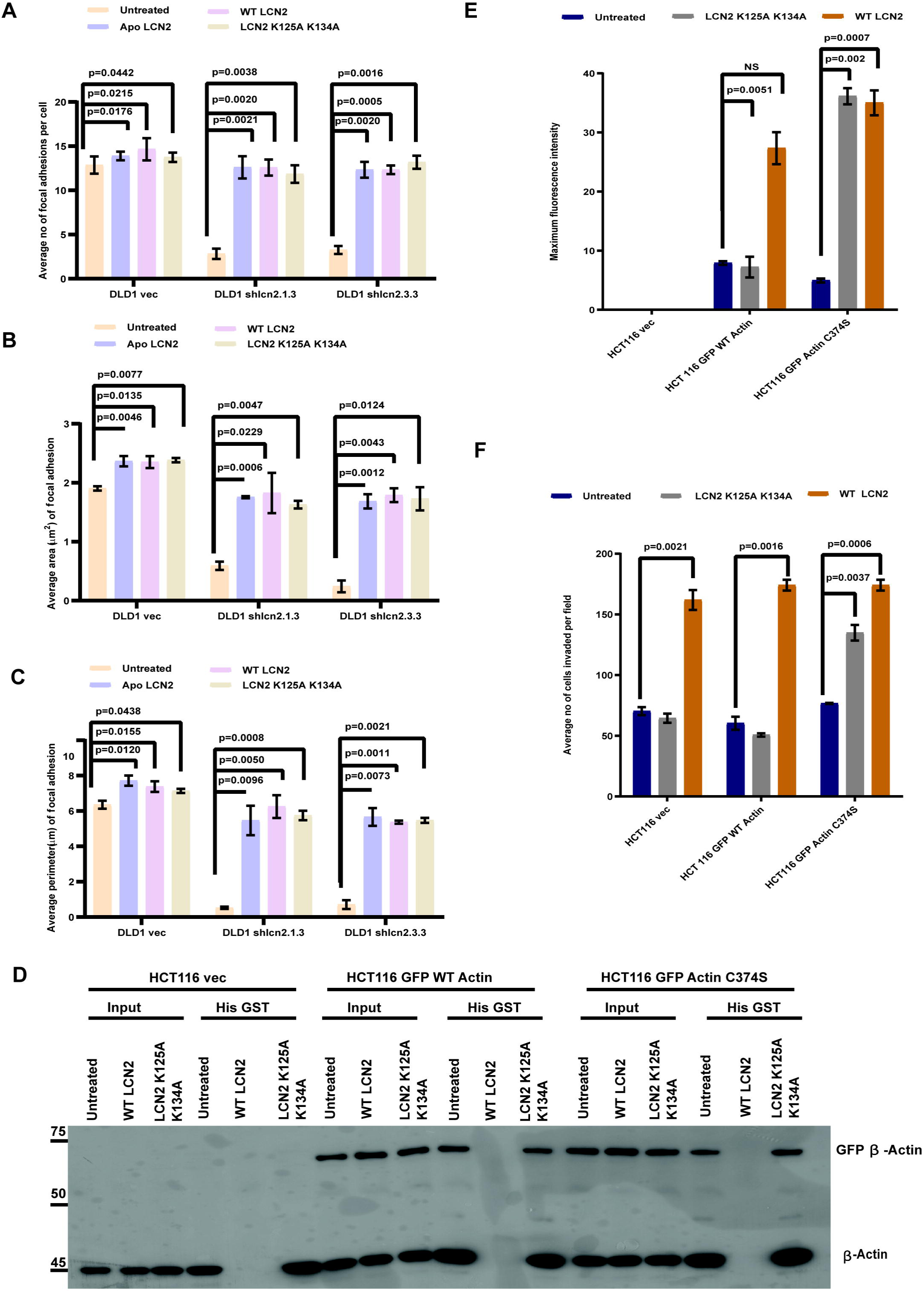
The ability of LCN2 to bind iron is not required for focal adhesion formation. **A-C.** The DLD1-derived LCN2 knockdown lines or the vector control were incubated with either the vehicle control (UT), WTLCN2, ApoLCN2 (not bound to iron), and the iron binding defective LCN2 mutant. Focal adhesion number (A), area (B), and perimeter (C) were determined in three independent experiments and the mean and standard deviation were plotted. **D-F.** Cells transfected with either the vector control, GFP-WT Actin, or GFP Actin C374S were incubated with either WT or mutant LCN2 produced in bacteria. Protein extracts were prepared and incubated with His GST or His LCN2 and the reactions resolved on SDS-PAGE gels followed by Western blotting with antibodies to actin (D). Cells were fixed and stained with TRITC-phalloidin and the levels of filamentous actin were quantitated in three independent experiments. The mean and standard deviation are plotted (E). Matrigel invasion assays were performed on the treated cells and the number of invading cells was determined in three independent experiments and the mean and standard deviation were plotted (F). Note that the presence of the C374S mutant results in an increase in filamentous actin formation and invasion in cells treated with iron-binding defective LCN2. Where indicated, p values were determined using a student’s t-test.

To identify the mechanisms underlying the increased focal adhesion formation stimulated by LCN2, we determined the levels of activated FAK and paxillin in LCN2 over-expressing or knockdown clones. Paxillin is phosphorylated at a Tyrosine residue at 118 (Y118) resulting in increased formation of nascent focal adhesions (Zaidel-Bar et al., 2007) and FAK is activated by auto-phosphorylation at Tyrosine 397 leading to the formation of a docking site for c-Src (reviewed in (Tilghman and Parsons, 2008)). c-Src phosphorylation of FAK leads to further FAK activation and phosphorylation of Paxillin at Y118 (Thomas et al., 1999). A Western blot analysis demonstrated that while Paxillin phosphorylation at Y118 was significantly elevated in LCN2 overexpressing cells and decreased in LCN2 knockdown cells, no changes were observed in the levels of FAK phosphorylated at Y397 (Fig. 3A-F). These results suggested that LCN2 might be stimulating the activation of c-Src. c-Src kinase activity is inhibited by phosphorylation at Tyrosine 527 (Y527) and stimulated by phosphorylation at Tyrosine 416 (Y416) (Okada, 2012; Xu et al., 1999b). A Western blot analysis demonstrated that the phosphorylation of c-Src at Y527 was significantly decreased and phosphorylation at Y416 increased in the LCN2 overexpressing cells (Fig. 3G-I). In contrast, LCN2 knockdown led to an increase in phosphorylation at Y527 and a decrease in phosphorylation at Y416 (Fig. 3J-L). Consistent with these data, treatment with an inhibitor of c-Src led to a significant decrease in Y118 phosphorylation on Paxillin and Y416 phosphorylation on c-Src in LCN2 overexpressing cells (Fig. 4A-C). The decrease in c-Src and Paxillin phosphorylation was accompanied by a decrease in the number, area, and perimeter of the focal adhesions in LCN2 overexpressing cells (Fig. 4D-F and SF4A) and a decrease in invasion (SF4B-C). Further, treatment with the anti-LCN2 antibody led to a significant decrease in Paxillin Y118 and c-Src Y416 phosphorylation and an increase in Y527 phosphorylation as compared to untreated cells or cells treated with MIgG (Fig. 4G-J). Taken together, these results indicated that LCN2 stimulates focal adhesion formation by stimulating the kinase activity of c-Src.

**Figure 3.**
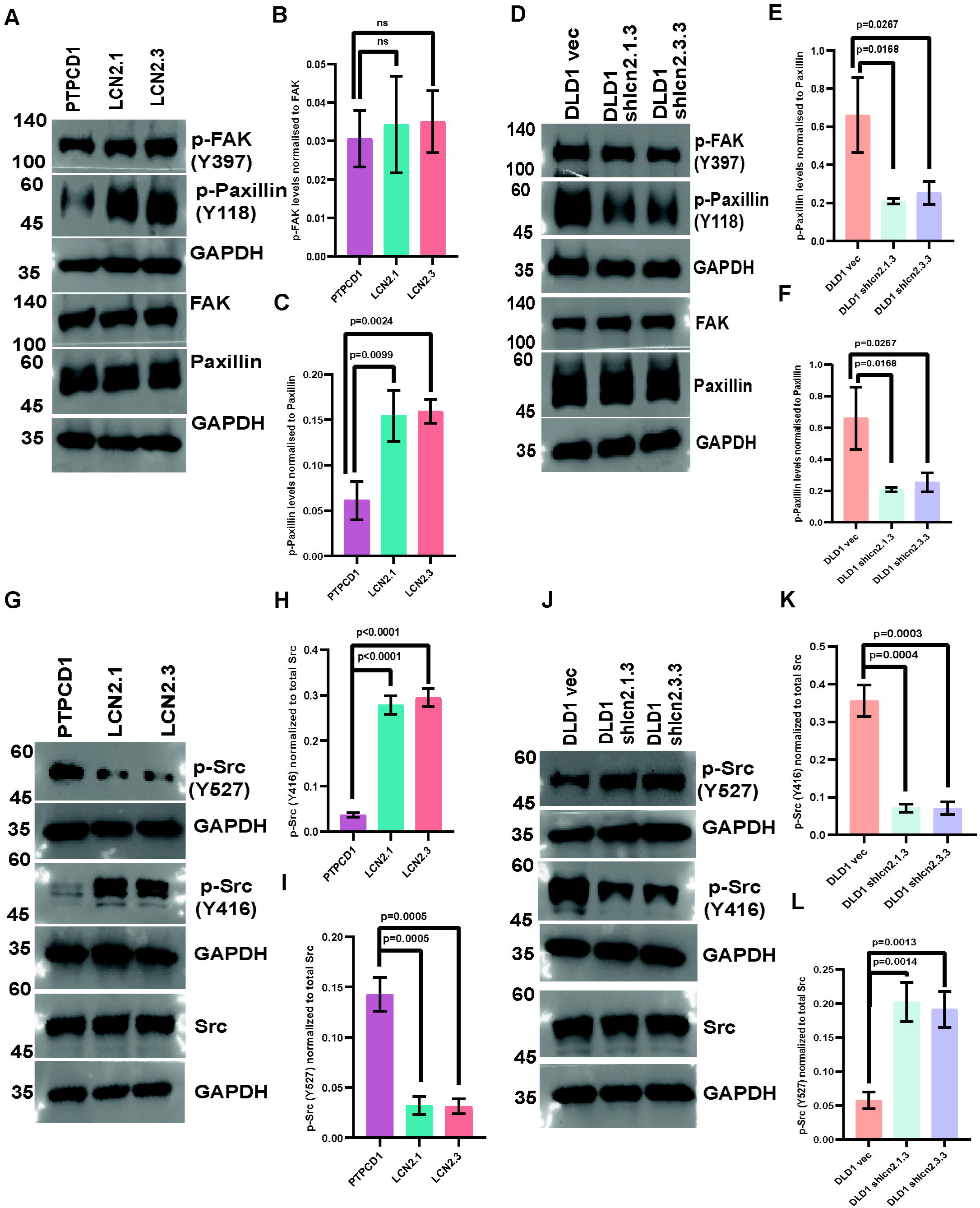
LCN2 expression stimulates c-Src activation leading to paxillin phosphorylation. **A-L.** Protein extracts prepared from the indicated cell lines were resolved on SDS-PAGE gels followed by Western blotting with the indicated antibodies. The intensity of the bands was determined using a BioRad chemidoc and the mean and standard deviation of three independent experiments is plotted. Note that the levels of p-FAK(Y397) are unaltered in cells with either elevated or low levels of LCN2 (A, B, D, E) while the levels of p-Paxillin (Y118) are elevated in LCN2 high cells and low in LCN2 knockdown cells (A, C, D, F). The levels of the activated c-Src (p-Src Y416) are elevated and lowered in LCN2 overexpressing and knockdown clones respectively (G, I, J, L), and the levels of inactive c-Src (p-Src Y527) are lowered and elevated in LCN2 overexpressing and knockdown clones respectively (G, H, J, K). Note that total levels of paxillin, FAK, and c-Src remain unchanged. Western blots for GAPDH served as loading controls for normalization. Where indicated, p values were determined using a student’s t-test.

**Figure 4.**
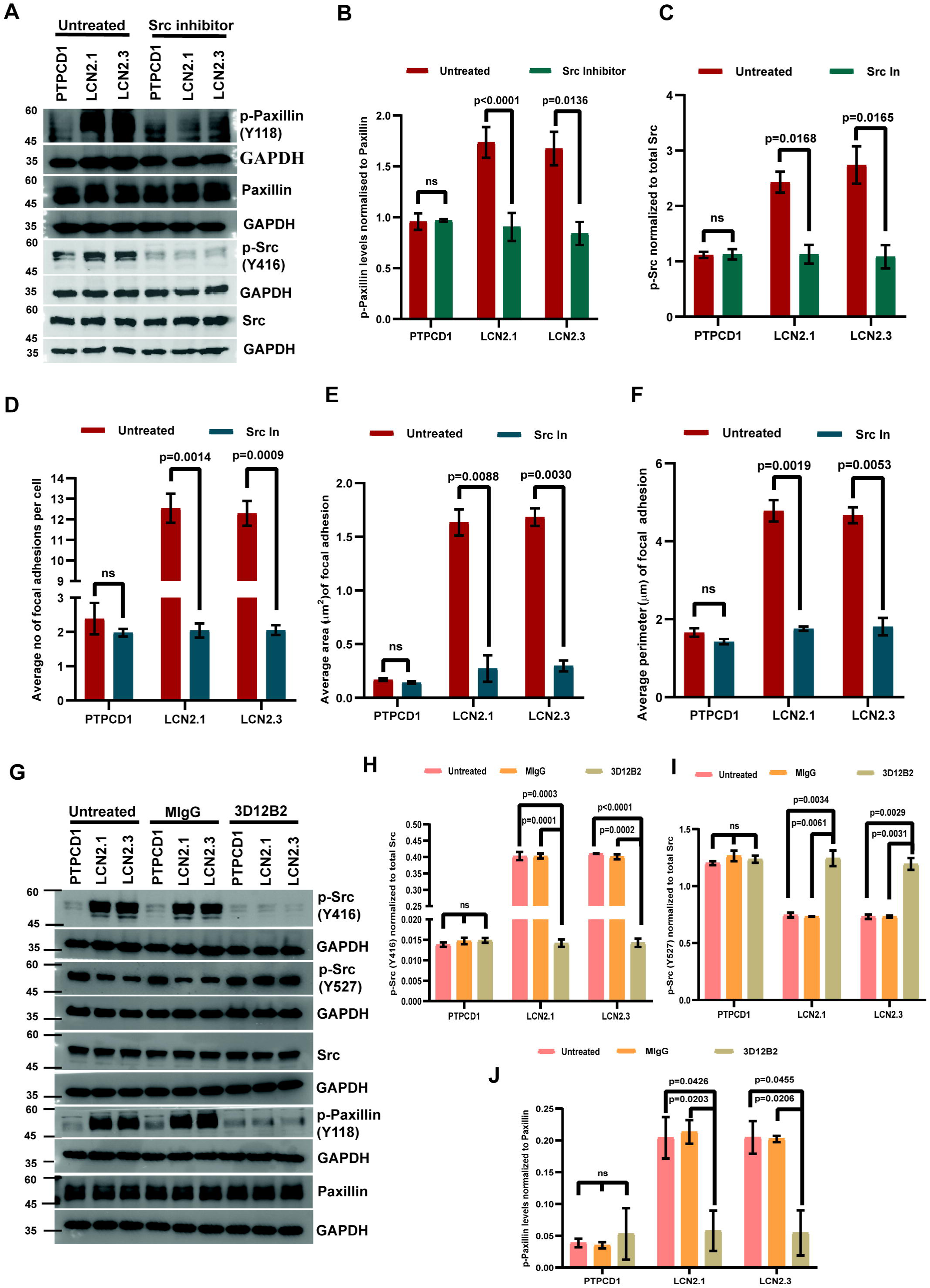
LCN2-mediated activation of c-Src leads to focal adhesion formation. **A-C.** Protein extracts prepared from the indicated cell lines that were untreated or treated with the c-Src inhibitor were resolved on SDS-PAGE gels followed by Western blotting with the indicated antibodies. The intensity of the bands was determined using a BioRad Chemidoc and the mean and standard deviation of three independent experiments are plotted. **D-F.** The indicated cell lines were stained with antibodies to paxillin or TRITC-phalloidin and imaged by confocal microscopy. The number (D), area (E), and perimeter (F) of focal adhesions were quantitated in three independent experiments and the mean and standard deviation were plotted. **G-J.** Protein extracts from cell lines that were untreated or treated with non-specific mouse IgG or the LCN2 antibody were resolved on SDS-PAGE gels followed by Western blotting with the indicated antibodies (G). The intensity of the bands was determined using a BioRad Chemidoc and the mean and standard deviation of three independent experiments are plotted (H-J). Note that treatment with the LCN2 antibodies results in a decrease in the activation of c-Src and paxillin formation. Blots for GAPDH serve as loading controls. Where indicated, p values were determined using a student’s t-test.

c-Src is activated when Y527 is dephosphorylated by the Tyrosine phosphatase, PTP1B (Zhu et al., 2007). PTP1B protein and mRNA levels were elevated in LCN2 overexpressing cells (Fig. 5A, B, and E) and decreased in LCN2 knockdown cells (Fig. 5C, D, and F). Similarly, treatment with the anti-LCN2 antibody led to a decrease in the protein levels of PTP1B as compared to untreated cells or cells treated with MIgG (SF5A-B). To determine if the increase in PTP1B levels in LCN2 high cells was responsible for increased c-Src activation, we generated knockdown lines for PTP1B (gene name PTPN1) (SF5C-D) and demonstrated that focal adhesion number, area, and perimeter were decreased upon PTP1B knockdown in LCN2 overexpressing cells as compared to the vector control (SF5E and Fig. 5G-I). The decrease in focal adhesion formation was accompanied by a decrease in invasion (SF5F and Fig. 5J), a significant decrease in the phosphorylation of Y416 and Y118 in c-Src and Paxillin respectively, and an increase in the phosphorylation of Y527 in c-Src (Fig. 5K-N). These results suggest that LCN2 activates c-Src by promoting the expression of PTP1B.

**Figure 5.**
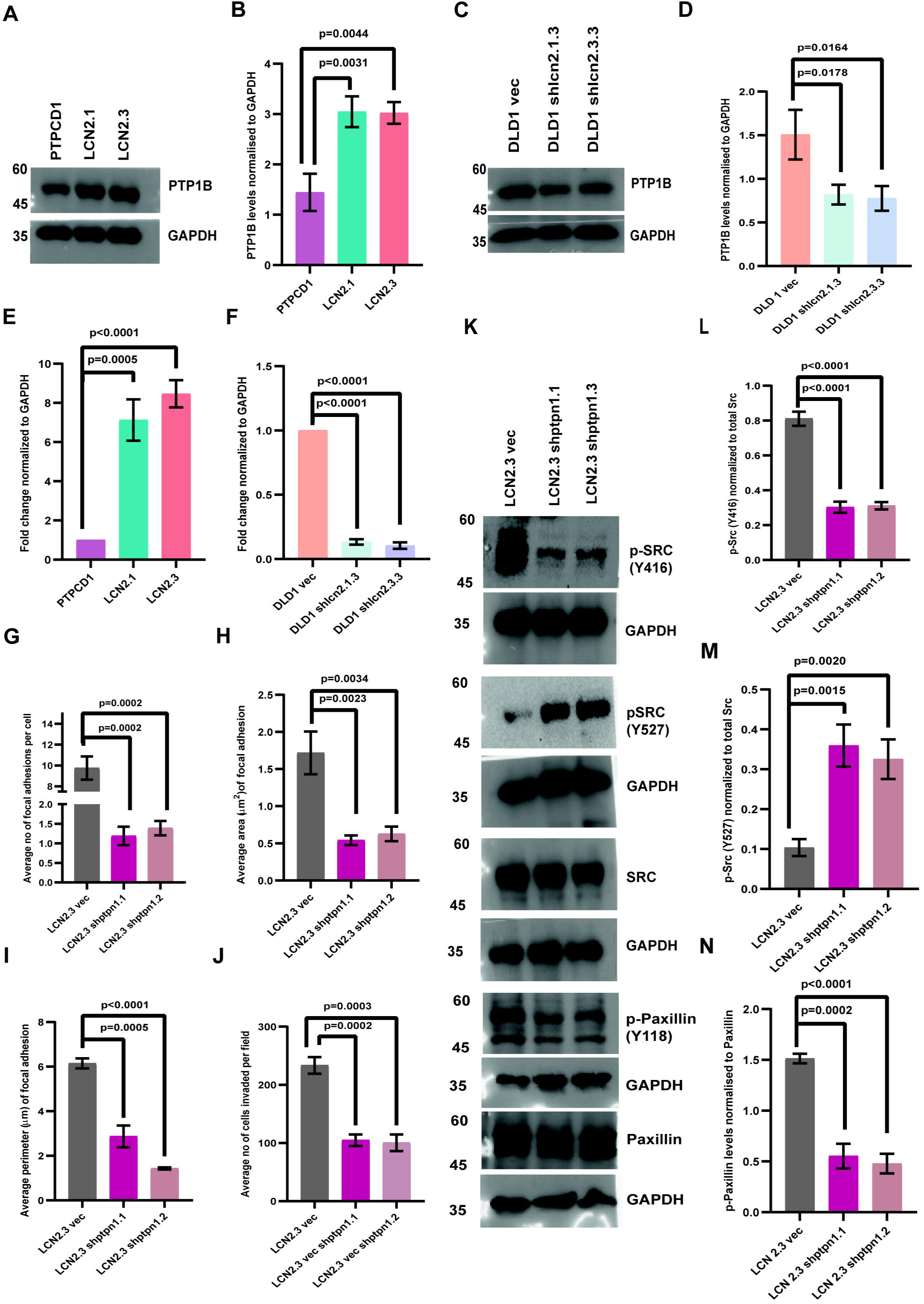
LCN2 expression leads to an increase in PTP1B levels and c-Src activation. **A-D.** Protein extracts from the indicated cell lines were resolved on SDS-PAGE gels and Western blots performed with the indicated antibodies (A&C). The mean and standard deviation were plotted for three independent experiments (B&D). E-F. mRNA prepared from the indicated cell lines was used as a template in reverse transcriptase-coupled quantitative PCR reactions. The mean and standard deviation for three independent experiments is plotted. **G-J.** The LCN2.3 derived vector control (LCN2.3vec) or PTP1B knockdown clones (LCN2.3shptpn1.1 and shptpn1.2) were stained with antibodies to paxillin and TRITC-phalloidin and the number (G), area (H) and perimeter (I) of focal adhesions was determined in three independent experiments and the mean and standard deviation plotted. The indicated cell lines were also used in invasion assays and the mean and standard deviation of the number of invading cells were determined in three independent experiments and the mean and standard deviation plotted (J). **K-N.** Protein extracts prepared from the indicated cells were resolved on SDS-PAGE gels followed by Western blotting with the indicated antibodies (K). The mean and standard deviation of three independent experiments were plotted (L-N). Western blots for GAPDH served as a loading control. Where indicated, p values were determined using a student’s t-test.

We have previously reported that LCN2 stimulates the expression of the transcription factor ETS1 (Chaudhary et al., 2021). The LCN2 overexpressing cells showed an increase in ETS1 protein and mRNA levels while the LCN2 knockdown cells showed a decrease in ETS1 protein and mRNA levels (SF6A-D). Similarly, inhibiting LCN2 function with the anti-LCN2 antibody led to a significant decrease in ETS1 levels as compared to the untreated cells or cells treated with MIgG (SF6E-F). A Jasper analysis demonstrated that the promoter of the PTP1B gene (PTPN1) contained binding sites for ETS1 (Fig. 6A). A correlation was observed between the expression of PTPN1 and ETS1 in colon adenocarcinoma samples (Fig. 6B). To confirm the contribution of ETS1 to PTP1B expression, we inhibited ETS1 expression in the LCN2 overexpressing cells using previously described shRNA constructs for ETS1 (Chaudhary et al., 2021) (SF6G-H). Loss of ETS1 led to a significant decrease in the levels of PTP1B protein and mRNA, and in c-Src and Paxillin phosphorylation at Y416 and Y118 respectively, which was accompanied by an increase in the levels of Y527 phosphorylation on c-Src (Fig. 6C-G). This was accompanied by a significant decrease in the number, area, and perimeter of focal adhesions (Fig. 6H-J) leading to decreased invasion as compared to the vector control cells (SF6I-J). Finally, chromatin immunoprecipitation assays demonstrated significantly increased occupancy of the PTPN1 promoter by ETS1 in LCN2 overexpressing cells as compared to the vector controls (Fig. 6K). These results suggest that LCN2 expression leads to an increase in c-Src activation by stimulating the expression of ETS1 leading to increased expression of PTP1B.

**Figure 6.**
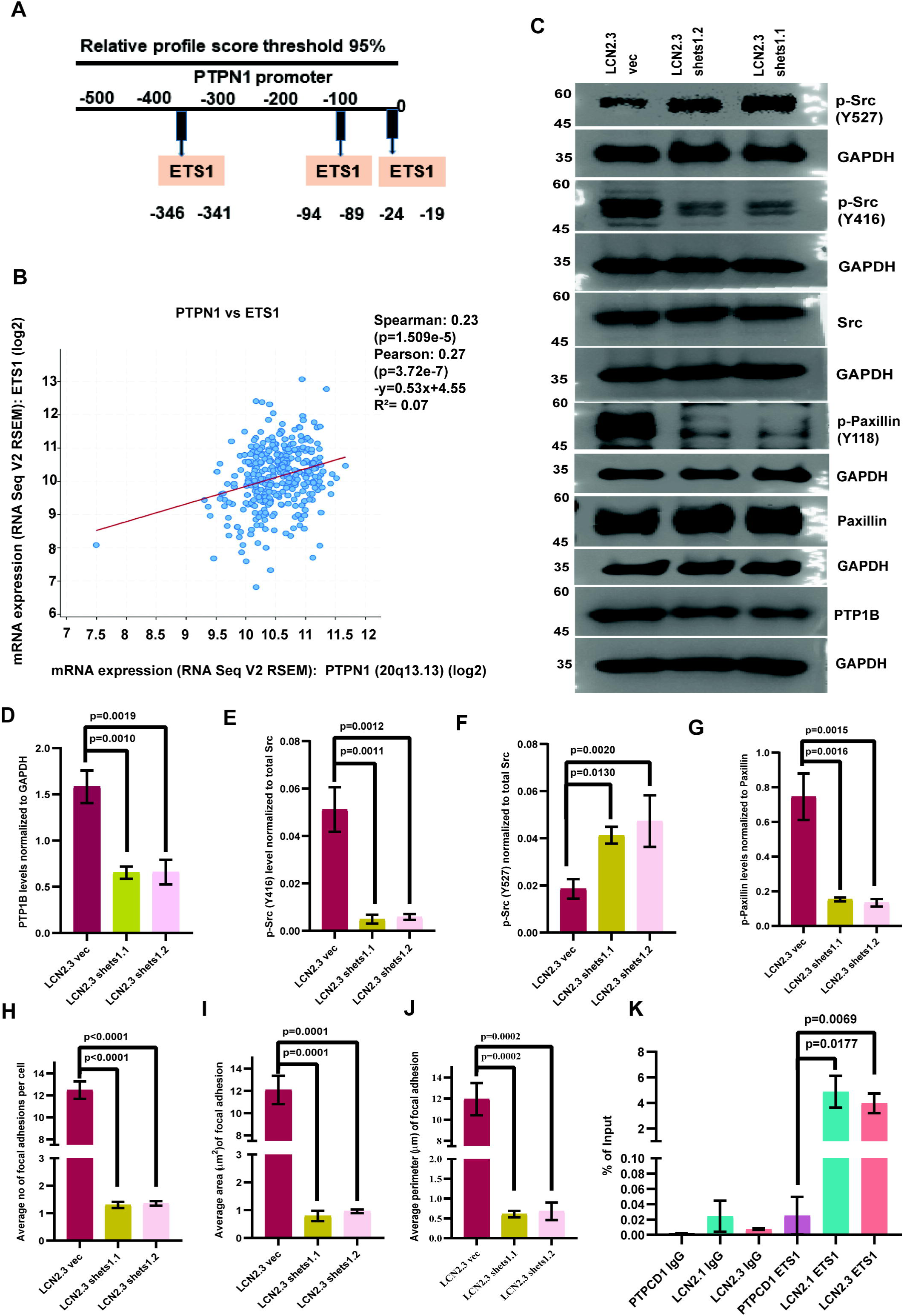
ETS1 expression is required for the PTP1B-mediated activation of c-Src. **A.** Jasper analysis of the PTP1B (gene name PTPN1) promoter identifies ETS1 binding sites in the promoter. B. Pearson’s coefficient analysis of PTPNI and ETS1 expression in the TCGA data set. The data are represented as a scatter plot where r and p refer to the correlation coefficient and p-value, respectively. **C-G.** Protein extracts prepared from the LCN2.3 derived vector control (LCN2.3vec) or ETS1 knockdown clones (LCN2.3ets1.1 and shets1.2) were resolved on SDS-PAGE gels followed by Western blotting with the indicated antibodies (C). The band intensity was determined in three independent experiments and the mean and standard deviation are plotted (D-G). Western blots for GAPDH serve as loading controls. **H-J.** The LCN2.3 derived vector control (LCN2.3vec) or ETS1 knockdown clones (LCN2.3ets1.1 and shets1.2) were stained with antibodies to paxillin and TRITC-phalloidin and the number (H), area (I) and perimeter (J) of focal adhesions determined in three independent experiments. The mean and standard deviation are plotted. **K.** ChIP assays were performed using either a non-specific rabbit IgG (IgG) or ETS1 antibodies from the indicated cell, followed by qPCR to detect the LCN2 promoter. The mean and standard deviation are plotted. Where indicated, p values were determined using a student’s t-test.

## Discussion

The data shown here demonstrate that LCN2 promotes invasion in two ways: the first by preventing actin glutathionylation allowing actin polymerization and the second by promoting focal adhesion formation. The first is dependent on the ability of LCN2 to sequester iron, thereby reducing ROS levels, and the second is dependent on the ability of LCN2 to promote the activity of c-Src leading to focal adhesion formation. Both LCN2 functions are required for the invasion of colon cancer cells.

Previous work from multiple laboratories has demonstrated that LCN2 promotes invasion and metastasis in multiple tumor types (Basu et al., 2018; Ding et al., 2015; Guo et al., 2014; Leng et al., 2009; Miyamoto et al., 2016; Oren et al., 2016; Yang et al., 2009). Further, Sun et al. demonstrated that increased LCN2 levels correlated with poor survival in CRC patients at all stages and that increased LCN2 levels correlated with increased invasion in the cecum and increased metastasis in mouse xenograft models (Sun et al., 2011). The ability of LCN2 to promote invasion has been ascribed to the ability of LCN2 to promote EMT (Ding et al., 2015; Yang et al., 2009), and to promote angiogenesis (Yang et al., 2013), suggesting that LCN2 could be a potential therapeutic target in multiple tumor types (Leng et al., 2009; Miyamoto et al., 2016). Indeed, potential therapeutic agents that inhibit LCN2 function show anti-tumor potential and inhibit invasion (Chaudhary et al., 2021; Choudhary et al., 2023; Guo et al., 2016; Guo et al., 2014; Leng et al., 2009; Yao et al., 2021). LCN2 levels can be stimulated by increased Her 2 activity (Leng et al., 2009), and an increase in p38 activity (Basu et al., 2018), and both lead to increased invasion and tumor formation. The results in this paper demonstrate that in addition to stimulating the polymerization of actin by inhibiting actin glutathionylation, LCN2 promotes the formation of focal adhesions by stimulating c-Src activity leading to increased invasion and migration.

Excessive iron leads to increased tumor progression in human colorectal cancer (Nelson, 2001) and in mouse models of colorectal cancer (Radulescu et al., 2012). Increased iron levels lead to the generation of ROS (Toyokuni, 1996), which is associated with colorectal cancer progression (Myant et al., 2013; Wang et al., 2017). However, high ROS levels inhibit the polymerization of G-actin, and high levels of ROS lead to the cleavage of actin filaments (Xu et al., 2017). Further, ROS can induce the glutathionylation of Cysteine 374 in actin leading to a decrease in filament stability and decreased polymerization (Stournaras et al., 1990; Wang et al., 2001). C374 glutathionylation is exacerbated in the presence of iron in the neurological disorder Freidrich’s Ataxia, leading to a decrease in migration (Pastore et al., 2003). All of these results suggest that increased ROS levels inhibit actin polymerization and invasion. However, multiple pieces of evidence suggest that invasion can be stimulated by the accumulation of ROS at the invasive front in tumor samples and in cells grown in vitro (Gianni et al., 2011; Gianni et al., 2010; Giannoni et al., 2005; Gozin et al., 1998). Increased ROS is associated with the increased formation of invadopodia (Gianni et al., 2010) and increased activation of FAK and c-Src (Giannoni et al., 2005; Gozin et al., 1998). These differences could be ascribed to the local accumulation of specific ROS species in cellular compartments and also due to the differences in total ROS (Cheung and Vousden, 2022; Tochhawng et al., 2013) and labile iron levels (Toyokuni, 1996). The results in this report indicate that in a high iron environment such as the colon, tumor cell invasion might be dependent on the ability of the tumor cell to clear ROS species by decreasing iron levels, thereby leading to a decrease in actin glutathionylation and increased actin polymerization.

Our results suggest that the actin mutant C374S, while not showing a great decrease in glutathionylation, shows increased polymerization and supports an increase in invasion in cells treated with the LCN2 K125AK134A mutant which is defective for iron binding. The discrepancy between glutathionylation of the C374S mutant could be that other Cysteine residues in actin are also glutathionylated as reported (Wilson et al., 2016), and that there might be a further increase in the glutathionylation of other residues in the C374S mutant. However, as previously reported, glutathionylation of C374 is the major modification that inhibits actin polymerization (Stournaras et al., 1990), which is consistent with the findings in this report. Our results also suggest that while the ability of LCN2 to bind iron is required for invasion and the inhibition of actin glutathionylation (Choudhary et al., 2023), iron binding by LCN2 is dispensable for the formation of focal adhesions. This is important as this is the first iron-independent function of LCN2 that has been reported and it opens up the possibility that there are other LCN2 functions required for tumor formation that are independent of the ability of LCN2 to regulate iron homeostasis.

Previous reports have suggested that c-Src phosphorylates ETS-1 resulting in increased stabilization of ETS-1 as it is not degraded by the E3 ligase COP1 leading to increased tumor progression (Lu et al., 2014). Further, ETS-1 is required for resistance to sorafenib in hepatocellular carcinoma by disrupting mitochondrial ROS (Vishnoi et al., 2022), leading to increased invasion. These data are consistent with our results which suggest that LCN2 promotes the activation of c-Src by stimulating the expression of ETS-1 leading to increased expression of the protein phosphatase, PTP1B. PTP1B dephosphorylates c-Src at Y527 resulting in increased activation of c-Src leading to increased paxillin phosphorylation, focal adhesion formation, and increased invasion. Inhibition of c-Src leads to a decrease in paxillin phosphorylation, focal adhesion number, and invasion. Further, we have previously reported that the increase in ETS-1 levels stimulated by LCN2 expression is required for the clearance of ROS (Chaudhary et al., 2021) and that ROS clearance by LCN2 is required to prevent actin glutathionylation and promote actin polymerization (Choudhary et al., 2023). Thus, in addition to promoting the formation of an active c-Src-FAK complex (Parsons et al., 2008) leading to focal adhesion formation, LCN2 also promotes actin polymerization by decreasing the levels of ROS.

Based on the results in this manuscript we propose the following model (Fig. 7). LCN2 promotes invasion in two ways: the first is by stimulating actin polymerization by preventing the glutathionylation of G-actin, which requires the ability of LCN2 to bind iron and inhibit ROS generation. LCN2 stimulates the expression of ETS-1 leading to c-Src activation and focal adhesion formation in a manner that does not require the ability of LCN2 to bind iron. This is the first report of an LCN2 function that is not dependent on the ability of LCN2 to bind iron. Both functions of LCN2 are required to promote invasion and LCN2 expression is associated with increased tumor stage in colorectal cancer (Chaudhary et al., 2021) and LCN2 is expressed in 60-70% of colorectal cancers (Chaudhary et al., 2021; Zhang et al., 2009). Thus, combining therapeutics targeting LCN2 and c-Src in invasive tumors might result in better patient outcomes and improved responses to chemotherapy in colorectal cancer.

**Figure 7.**
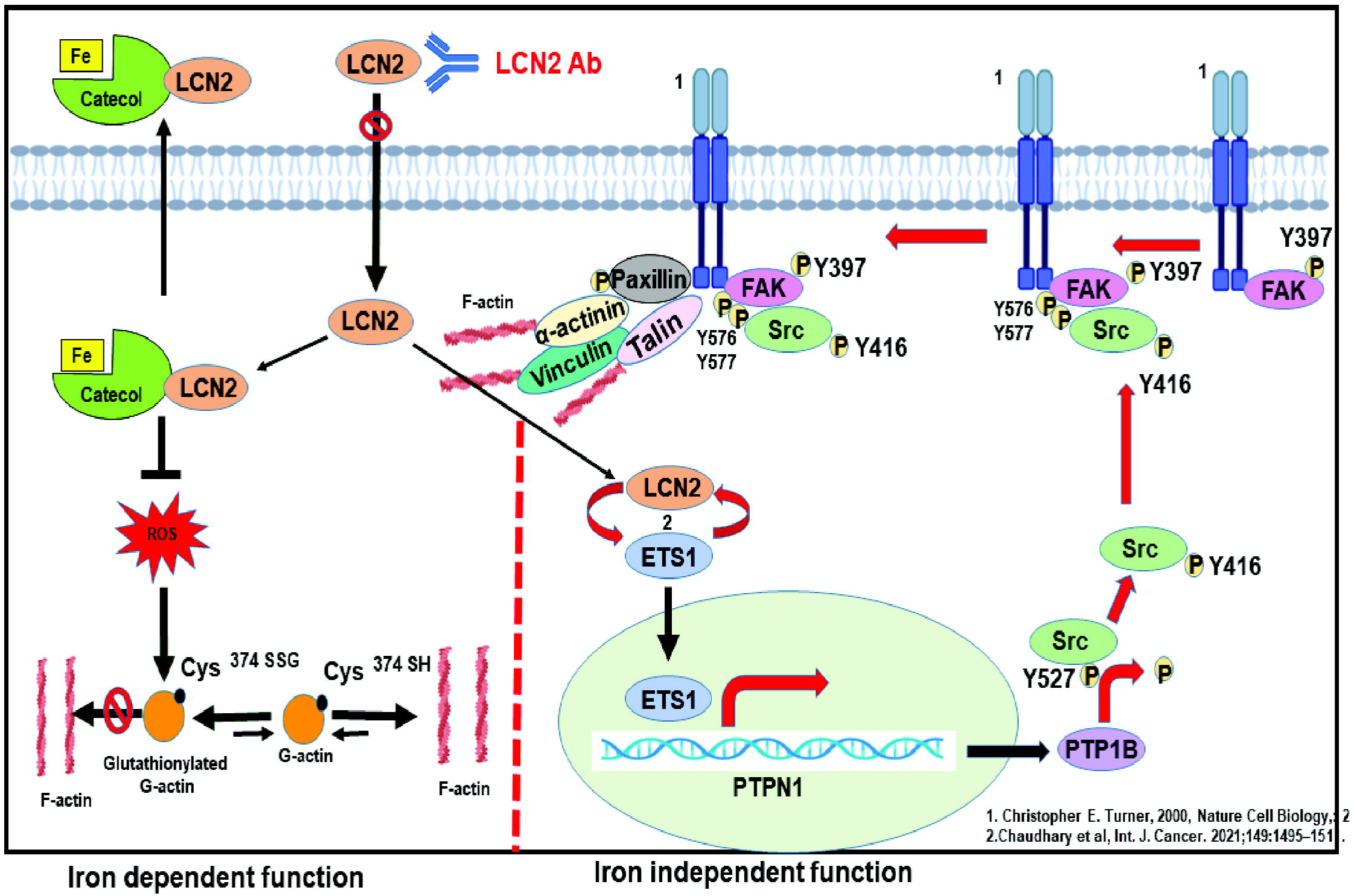
Model for how LCN2 stimulates invasion.

## Materials and Methods

### Cell lines and transfections

HCT116 cells (RRID: CVCL_0291) were procured from Dr. Bert Vogelstein, and DLD-1 (RRID: CVCL_0248) were procured from Dr. Sanjeev Galande and cultured as described (Choudhary et al., 2023). Cell lines were authenticated by short tandem repeat (STR) profiling during the last three years. Mycoplasma-free cells were used for all the experiments. The HCT116-derived LCN2 overexpressing lines (LCN2.1 and LCN2.3) and the vector control (PTPCD1) derived from HCT116 the DLD1-derived LCN2 knockdown clones (DLD1 shlcn2.1.3 and DLD1 shlcn2.3.3) and vector control (DLD1 vec) cells were previously described (Choudhary et al., 2023). The vector control and PTP1B knockdown clones were generated in LCN2.3 by transfecting these cells with an shRNA construct against PTP1B and maintained in 0.5μg/ml puromycin (Sigma Catalog #8833) ETS1 knockdowns were performed as described (Chaudhary et al., 2021). To determine the role of c-Src in invasion, migration, and focal adhesion formation, LCN2 overexpressing cells were treated with c-Src inhibitor (Sigma Catalog #S-2075)or vehicle control (DMSO) for 24 hrs.

### Plasmids

The shRNA constructs for LCN2 and ETS1 have been described previously (Chaudhary et al., 2021). The oligonucleotide sequences used to generate the PTP1B shRNA are in Supplementary Table 1. The oligonucleotides used to generate the C374S mutant are in Supplementary Table 1.

### Antibodiesand Western blot analysis

Whole-cell protein extracts were made either in EBC lysis buffer or in 1X sample buffer as described (Basu et al., 2015). Proteins from the cell supernatant were precipitated in acetone and lysate was prepared in 1X sample buffer for Western blots (Chaudhary et al., 2021). Dilutions of antibodies are shown in Supplementary Table 2. The blots were developed using either Super Signal West Pico Chemiluminescent Substrate (Pierce), Super Signal West Femto Chemiluminescent Substrate (Pierce), and Clarity Western ECL substrate (Bio-Rad) as per the manufacturer’s instructions. The anti-LCN2 antibody, 3D12B2, was generated and purified as described in (Chaudhary et al., 2021).

### Immunofluorescence and invasion assays

Immunofluorescence was performed as described previously (Basu et al., 2018). Dilutions used for primary and secondary antibodies are in Supplementary Table 3. Confocal images were obtained using an LSM780 Carl Zeiss confocal microscope on 63X objective with 2X optical zoom. The quantitation of paxillin streaks was performed using Fiji software. Paxillin streaks were measured by quantitating points having maximum intensity from find maxima by setting the prominence at 80. The area and perimeter of the paxillin streak were measured by drawing an ROI around individual streaks. 30 cells were used to quantify the paxillin streak number, area, and perimeter in three independent experiments. Matrigel degradation assays were performed using Boyden chambers as described previously [2].

### Actin glutathionylation assays

HCT116 cells were transfected with 1µg of either pEGFP Actin vector (Clontech Catalog #6116-1) or pEGFP Actin C374S or pEGFP N1 vector (Clontech Catalog #6085-1) and were selected in 5ug/ml Neomycin (Sigma Catalog #A1720) for 10 days. Actin glutathionylation using the His GST protein was determined as described previously (Choudhary et al., 2023), and the maximum fluorescent intensity of actin filaments was measured with Fiji software.

### ChIP assays

Cell lines were grown to 60%-70% confluence in 10 cm culture dishes were used and fixed with 1% (v/v) formaldehyde for 10 min at room temperature followed by quenching with 0.25 M glycine. The cells were sonicated to lyse and fragmentation of chromatin for size (200-600 bp). 5% of the lysed chromatin served as input for control, and the rest of the lysate was equally divided to be used for immunoprecipitation with either non-specific IgG or the ETS1 antibody. The DNA eluted from immunoprecipitated lysate and proceeded for RT-PCR using primer against PTPN1. The oligonucleotides used for the chromatin immunoprecipitation experiments are in Supplementary Table 1. The percentage of pull-down chromatin was determined using the amount of input DNA.

### Bioinformatics Analysis

cBioPortal software (https://www.cbioportal.org) was used to determine the correlation between ETS1 and PTPN1 by retrieving the RNA seq data of 342 samples for colorectal adenocarcinoma from the TCGA dataset.

## Acknowledgments

The authors wish to acknowledge the ACTREC Digital Imaging Facility for help with generating images.

## Competing interests

There are no competing interests

## Funding

The experiments in this report were funded by a grant from the Department of Biotechnology (BT/PR44632/BRB/10/1999/2021), Department of Atomic Energy DPR Nos. 1/3(7)/2020/TMC/R&D-II/ 8823 and 1/3(6)/2020/TMC/R&D-II/ 3805 and donations to the ACTREC Basic Research fund (4338) to SND. NC is a DST Inspire faculty (award no. IFA22 LSBM 271).

## Data availability

All relevant data can be found within the article and its supplementary information.

## Notes

### Competing Interest Statement

The authors have declared no competing interest.

